# Quality control for the target decoy approach for peptide identification

**DOI:** 10.1101/2022.11.17.516857

**Authors:** Elke Debrie, Milan Malfait, Ralf Gabriels, Arthur Declerq, Adriaan Sticker, Lennart Martens, Lieven Clement

## Abstract

Reliable peptide identification is key in mass spectrometry (MS) based proteomics. To this end, the target-decoy approach (TDA) has become the cornerstone for extracting a set of reliable peptide-to-spectrum matches (PSMs) that will be used in downstream analysis. Indeed, TDA is now the default method to estimate the false discovery rate (FDR) for a given set of PSMs, and users typically view it as a universal solution for assessing the FDR in the peptide identification step. However, the TDA also relies on a minimal set of assumptions, which are typically never verified in practice. We argue that a violation of these assumptions can lead to poor FDR control, which can be detrimental to any downstream data analysis. We here therefore first clearly spell out these TDA assumptions, and introduce TargetDecoy, a Bioconductor package with all the necessary functionality to control the TDA quality and its underlying assumptions for a given set of PSMs.

**TOC Graphic:** 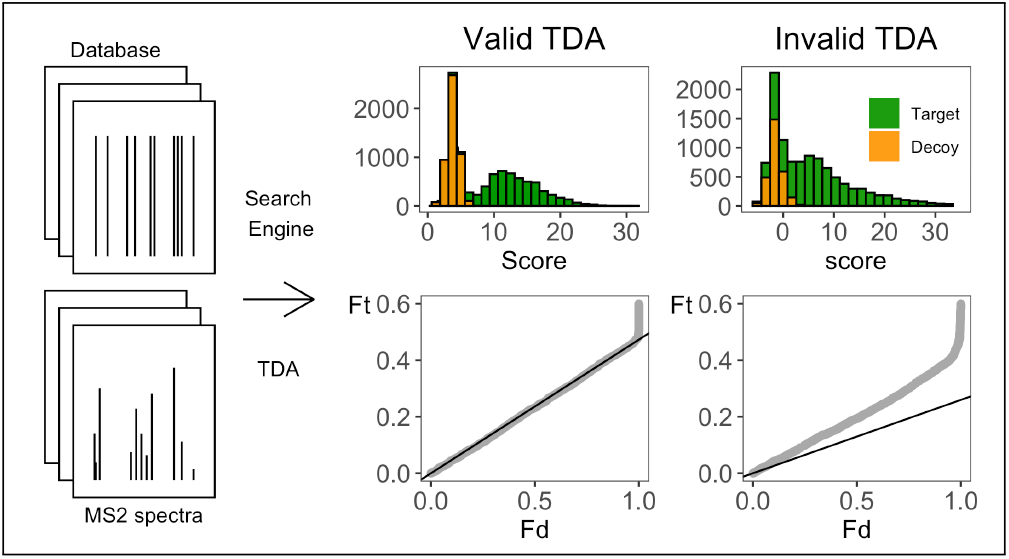

## Introduction

Peptide identification is an important step in the data analysis of mass spectrometry (MS) based proteomics^1^. In this step, search engines are typically used to match the large number of acquired experimental mass spectra to theoretical peptides derived from a sequence database. It is, of course, crucial for downstream analysis to evaluate the quality of these peptide-spectrum matches (PSMs). Therefore, False Discovery Rate (FDR) control is adopted to ensure the return of a reliable list of PSMs. The FDR estimation, however, requires good characterization of the score distribution of PSMs matched to a wrong peptide sequence (incorrect PSMs). In proteomics, the distribution of incorrect PSMs is estimated through the target decoy approach (TDA), which is typically viewed as a universal solution for assessing the FDR of a set of PSMs. The concatenated TDA method matches spectra to a database of real (targets) and nonsense peptides (decoys). A popular approach to generate these decoys is to reverse the target database. Hence, all PSMs that match a decoy sequence are known to be incorrect PSMs and the distribution of their scores is used to estimate the distribution of the scores of incorrect target PSMs. Note, that a crucial assumption of the TDA is therefore that decoy PSMs have similar properties to incorrect target PSMs, and that decoy PSM scores thus provide a good approximation of the distribution of incorrect target PSM scores. Users, however, typically do not evaluate these key assumptions, and blindly rely on the FDR returned by the TDA to select a set of mostly reliable PSMs for their downstream analysis.

We argue that mass spectrometrists and data analysts should critically verify the TDA assumptions to avoid incorrect peptide identifications to be labeled as reliably correct ones, which can have a detrimental impact on downstream data analysis. Indeed, Elias and Gygi, 2007, who developed the TDA, acknowledge that it imposes a few assumptions^2^, and, Gupta et al., 2011, amongst others, argued that some popular MS/MS search tools are not TDA-compliant^3^. Moreover, the TDA is also the default method in more challenging applications involving modified peptides, where it is as yet unknown how to construct reliable decoys, and where novel decoy strategies have been introduced without carefully evaluating their properties. Despite the resulting clear need, no tools have been developed so far to critically assess TDA assumptions, leaving mass-spectrometrists in the dark on the quality of the FDR that is associated with their list of, assumed reliable, PSMs.

We here therefore first carefully dissect the TDA method to unravel its underlying assumptions. We then present diagnostic plots to critically evaluate these assumptions, and we introduce TargetDecoy, our novel Bioconductor package with functions for quality control of the TDA and its underlying assumptions. Finally, we illustrate its use on data from different applications, showing that it provides the necessary functionality to assess the overall quality of search results.

## Methods

In the peptide identification step, search engines are typically used to match the large number of acquired experimental mass spectra to theoretical peptides derived from a sequence database. These peptide to spectrum matches (PSMs) can either match to the correct peptide sequence, which we refer to as a correct PSM, or to a wrong peptide sequence, which we refer to as an incorrect PSM. Therefore, False Discovery Rate (FDR) control is typically adopted to ensure that a reliable list of PSMs is returned. The competitive target decoy approach(TDA) is the default method that is used to estimate the FDR. However, the TDA does not come without assumptions that should be met. Here, we start by introducing some basic concepts. Then, we dissect the TDA method so as to unravel its underlying assumptions. Next, we introduce the datasets that will be used in this paper to illustrate our method and we conclude with the implementation of our tool. The diagnostic plots that we developed for the evaluation of the TDA assumptions are introduced in detail in the results section.

### Background

In the peptide identification step search engines return for each observed spectrum a peptide to spectrum match (PSM), i.e. the peptide sequence from the database that has the spectrum that is most similar to the observed spectrum according to the score metric of the search engine. So the PSM is the peptide sequence with the “best” score among all candidate peptide sequences in the database. Depending on the metric of the search engine, the “best” score is either the lowest or the highest one. Without loss of generality, however, we can develop diagnostic plots in which it is assumed that better scores are higher. Indeed, we can simply transform the score metric of the search engine, e.g. by changing its sign. Search engines often return e-values, i.e. probabilities to observe a random match in the database with a score that is higher than the one observed for a particular PSM. Hence, smaller evalues are better and the best e-value scores are all compressed in the region close to zero, which does not provide a good visual discrimination for PSMs with a reliable score. In our TargetDecoy tool we therefore provide an optional argument to transform e-values with a −log10 transformation, which effectively transforms e-values to a score for which higher values are better and thus allows for a better visual inspection of how well the observed spectrum of the candidate PSMs is matching to the theoretical spectrum according to the score metric of the search engine. In the remainder of the paper, higher scores indicate that the observed spectrum is matching better to the theoretical spectrum of its PSM, at least according to the search engine’s score metric.

### Assumptions of the competitive target decoy approach

The competitive TDA involves a search on a concatenated target and decoy sequence database. Typically, one will ensure that there are as many target as decoy sequences in the concatenated database. If one uses a competitive TDA with target and decoy databases of equal size and if decoy PSMs are a good approximation of incorrect target PSMs, then it is equally likely that an incorrect PSM will match to a wrong target sequence as to a decoy sequence.

Because decoy PSMs are incorrect, the TDA only has to control the FDR of the set of reliable target PSMs that is returned for downstream analysis. Note, the target PSM scores follow a mixture distribution:

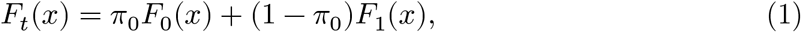

with *F_t_*(*x*) is the cumulative distribution of the scores of the target PSMs, *π*_0_ the fraction of incorrect target PSMs, *F*_0_(*x*’) the cumulative distribution of the scores of incorrect target PSMs and *F*_1_(*x*’) the cumulative distribution of the scores of correct target PSMs.

It is well known from the statistical literature^4^ that the FDR of a set of PSMs above a threshold *t* equals

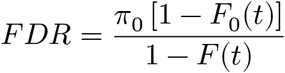

With the competitive TDA, the FDR of the set of returned PSMs with scores above a threshold *t,* is estimated by dividing the number of decoy PSMs with a score above *t* by the number of target PSMs with a score above *t*:

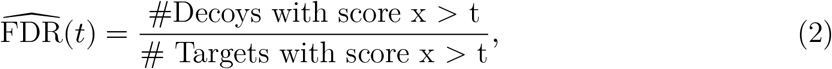

which we can rewrite as

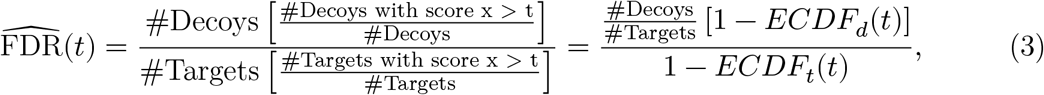

with *ECDF_d_*(*t*) and *ECDF_t_*(*t*) the empirical cumulative distribution function of decoys and targets, respectively. Hence, this result implies that the distribution of incorrect target PSMs, *F*_0_(*x*) is implicitly estimated by the ECDF of the decoys, i.e. 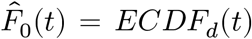 and the proportion of incorrect PSMs among the targets, 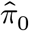 is estimated as the ratio of the number of decoy PSMs on the number of target PSMs. Hence, two assumptions have to be checked:

1. that decoy PSMs are a good approximation of incorrect target PSMs and that their distributions are thus (very close to) equal; and
2. that an incorrect PSM is equally likely to match to a target sequence as to a decoy sequence when the database of target and decoy sequences is equal in size.

### Datasets

MS runs from three publically available datasets were downloaded from the ProteomePro-teomeXchange Consortium (http://proteomecentral.proteomexchange.org) PRIDE partner repository^5^: A *Pyrococcus furiosus* dataset, a human dataset, and an immunopeptidomics dataset, with identifiers PXD001077, PXD028735, and PXD021398, respectively^6–8^. Raw MS datasets were fully reanalyzed, except for the immunopeptidomics dataset, where publically available search results were used. The “Velos005137” run was used from the *P. furiosus* dataset, and the “LFQ_Orbitrap_DDA_Human_01” run was used from the human dataset. Through a custom Nextflow workflow (Nextflow v22.10.0.5826) using Biocontainer images, raw MS files were converted to MGF with ThermoRawFileParser v1.4.0 and searched with the X!Tandem and MS-GF+ search engines through the unified SearchGUI command line interface v4.0.41^9–14^. Finally, search results were converted into a uniform tab-separated value format with psm_utils v0.1.1^15^.

Three distinct search spaces were used: (1) “Swiss-Prot Pyrococcus”, which includes all canonical Swiss-Prot protein sequences for *P. furiosus* (taxon ID 186497, downloaded on 03/11/2022, 504 entries), (2) “Swiss-Prot Human”, which includes all canonical Swiss-Prot protein sequences for *H. sapiens* (taxon ID 9606, downloaded on 03/11/2022, 20401 entries), and (3) “UniProt Human”, which includes all canonical and isoform protein sequences from the UniProt reference proteome of *H. sapiens* (taxon ID 9606, downloaded on 03/11/2022, 102572 entries). Each search space was complemented with the GPM cRAP contaminants database (downloaded from http://ftp.thegpm.org/fasta/cRAP/crap.fasta on 3/11/2022, 116 entries).

MS-GF+ and X!Tandem searches were performed with identical search settings, configured through SearchGUI. Carbamidomethylation of C was set as fixed modification, oxidation of M and phosphorylation of S, T, and Y were set as variable modifications. Precursor tolerance was set at 10 ppm and fragment tolerance at 0.5 Da. All other search settings were left as set by default in SearchGUI. For each search setup, X!Tandem was run twice: With the refinement option set to on or set to off.

The immunopeptidomics search results were used as is without any alterations from PXD021398. The “Figure_3_MaxQuant_100%FDR_and_Rescoring.zip” was downloaded from the PRIDE repository wherefrom the “msms_IAA.txt” file was extracted and was also converted into a uniform tab-separated value format with psm_utils v0.1.1.

The custom workflow, sequence databases, SearchGUI parameter files and search results are available on Zenodo at https://doi.org/10.5281/zenodo.7308022.

### Implementation

TargetDecoy is an open source R package available on Bioconductor (https://bioconductor.org/packages/release/bioc/html/TargetDecoy.html) and is released under the Artistic-2.0 license. It is a lightweight R package for checking the TDA assumptions in a single search, or in larger projects that involve multiple searches. It builds upon CRAN and Bioconductor R-packages and uses some code from the fdata-selection.R function from MSnBase^16^. R can be installed on any operating system from CRAN (https://cran.r-project.org/) after which you can install TargetDecoy by using the following commands in your R session:

~~~
if (!requireNamespace(“BiocManager”, quietly = TRUE)) {
    install.packages("BiocManager")
 }
BiocManager::install(“TargetDecoy”)
~~~

TargetDecoy currently supports objects of class tibble or data.frame, mzID or mzRIdent from the mzID and mzR packages, respectively. These datasets are typically loaded from an .mzid file as follows:

~~~
< filename <-“/path/to/file”
~~~

Using the mzID package

~~~
## mzID
> mzID_object <-mzID::mzID(filename)
~~~

by using the mzR package

~~~
## mzRIdent
> mzR_object <- mzR::openIDfile(filename)
~~~

or by loading a tsv file

~~~
> tibble_object <- read_tsv(filename)
~~~

An example dataset is available in the package. The results of an MS-GF+ search are stored in the ModSwiss object, which is an mzID object. It can be loaded through

~~~
> data(ModSwiss)
~~~

A histogram and percentile-percentile (P-P) plot of target and decoy scores for the different PSMs can be generated using the evalTargetDecoy function. It has the following arguments: ‘object’ is an mzID object, an mzR object or a data.frame object with the search results, ‘score’ is a character string with the name of the variable used to store the search engine scores, ‘decoy’ is a character string with the name of a Boolean variable indicating whether the PSM was matched to a decoy (TRUE) or to a target (FALSE), ‘log10’ is a Boolean variable indicating whether, scores are −log_{10} transformed or not, and ‘nBins’ is an integer that indicates the number of bins used to construct the histogram.

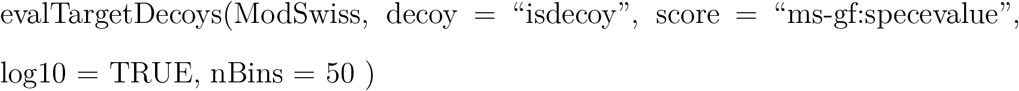

Sometimes variable names are not known upfront. If this is the case, the evalTargetDecoys() functions can be called with only an input object (and any variables that are already known). This launches a Shiny^17^ gadget that allows interactive selection of the variables. A histogram and P-P plot of the selected variables are created on the fly for previewing, together with a snapshot of the selected data (See Figure 1).

**Figure 1:**
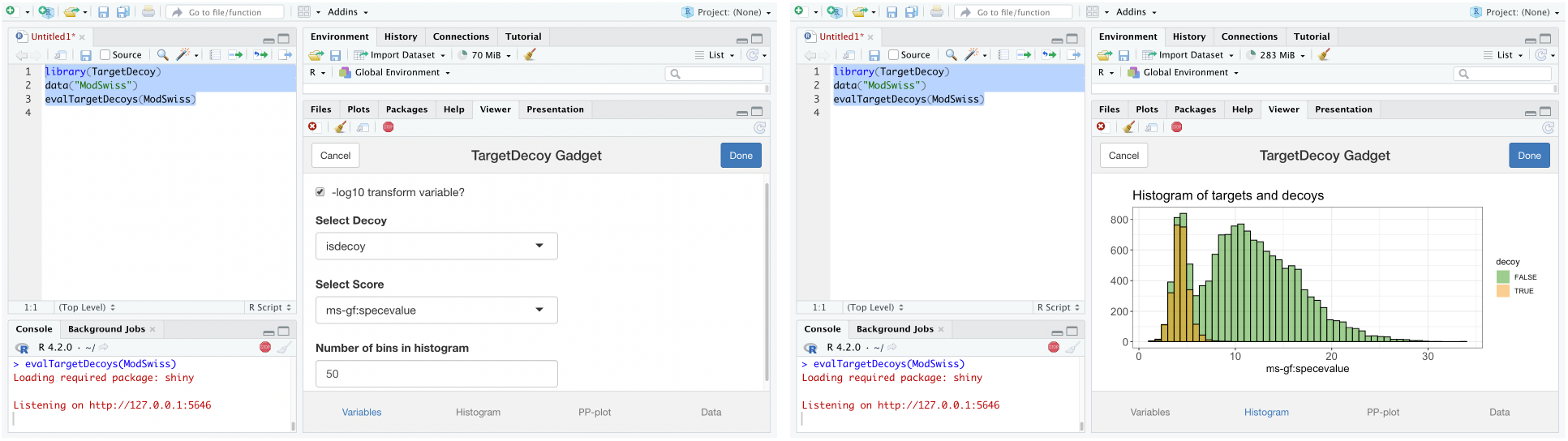
Screenshot of the Shiny gadget for interactive variable selection and plotting (here in R-studio)

When working in RStudio, the gadget opens in the “Viewer” pane by default, but it can also be opened in any browser window.

The code to generate all plots in this manuscript is available on our companion github page https://statomics.github.io/TargetDecoyPaper/.

## Results & Discussion

In this section we first illustrate the TDA method. Next, we develop diagnostic plots to evaluate the target decoy assumptions and we conclude with illustrating these in case studies on *Pyrococcus furiosus, Homo sapiens* and peptidomics PSM data.

A first and important step in an MSdata analysis workflow is peptide identification, where the large number of acquired experimental mass spectra are typically matched to theoretical peptides derived from a sequence database. Search engines return a peptide to spectrum match (PSM) that matches an observed spectrum to the peptide with the “best” score among all theoretical spectra of candidate peptide sequences in the database. The output of the search engine will contain correct PSMs that match to their proper peptide sequence, and incorrect PSMs that match to the wrong peptide sequence. The corresponding score distributions of correct and incorrect PSMs are typically separated, however, their tails overlap as we can also expect some correct PSMs to have a low score (see left panel of Figure 2).

**Figure 2:**
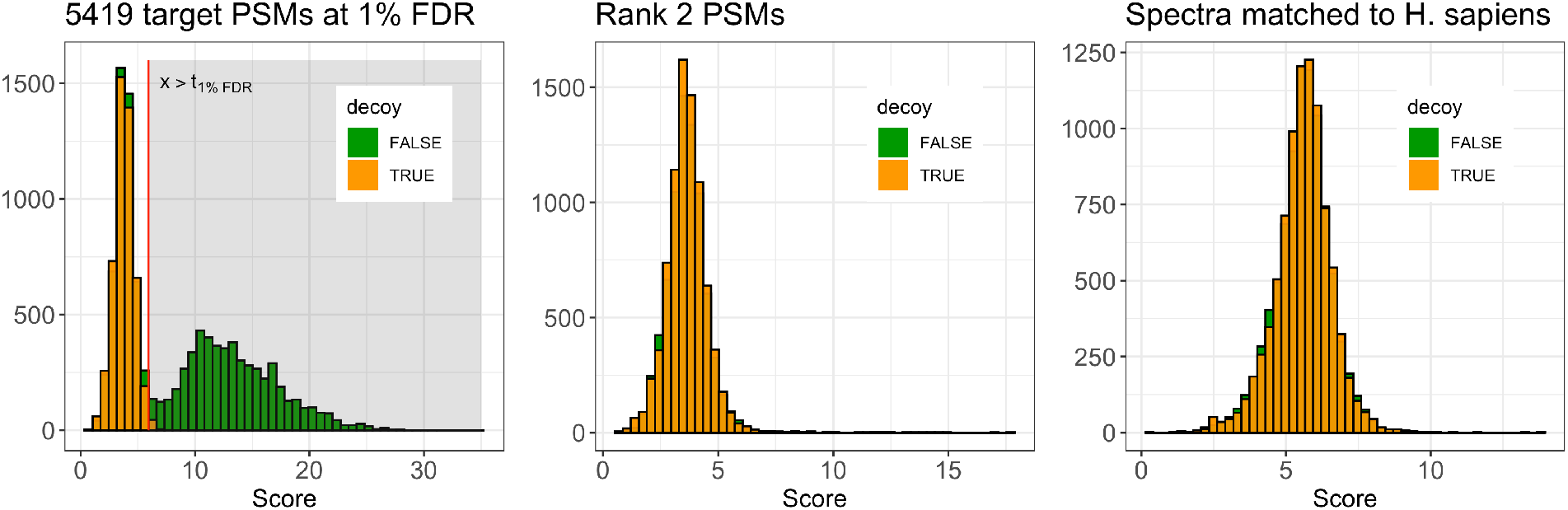
Histogram of targets and decoy scores for a concatenated search on a *Pyrococcus* run using MS-GF+. Left panel: Target and decoy PSM scores for an MS-GF+ search against the Swiss-Prot *Pyrococcus* database. The distribution of the target PSMs is bimodal. A higher score for −log10(ms-gf:specevalue) indicates that the theoretical spectrum of the PSM matches better with the observed spectrum according to the MS-GF+ scoring metric. Middle panel: Target and decoy scores for the rank 2 PSMs of the same search as in the left panel. MS-GF+ can also return the score for the second best PSM (rank 2), i.e. the peptide sequence with a theoretical spectrum that matches the observed spectrum with the second best score. Right panel: Target and decoy PSM scores for an MS-GF+ search against the Swiss-Prot Human database. None of the observed spectra is expected to have their proper match in the database.

In the Methods, we showed that the competitive TDA estimates the distribution of the scores of all target PSMs empirically, and that of incorrect target PSMs using the empirical distribution of the decoy PSMs. The left panel of Figure 2 shows histograms of the PSM scores for a *Pyrococcus furiosus* MS run searched with MS-GF+ against the Swiss-Prot *Pyrococcus* search space using the competitive TDA method. The distribution of the target PSMs is bimodal. A higher score for −log10(ms-gf:specevalue) indicates that the theoretical spectrum of the PSM matches better with the observed spectrum according to the MS-GF+ scoring metric. Hence, the left mode of the target distribution is likely to be enriched with incorrect target PSMs. The tails of the two distributions seem to overlap, indicating that we can also expect correct target PSMs with a low score. The distribution of the decoy PSM scores has the same shape as the first mode of the distribution of the target PSM scores. The height of the decoy histogram is a bit lower than the height of the first mode of the target distribution. Again, this can be expected because a few low scoring target PSMs are also likely to be correct matches. Note, however, that it is less evident to assess the decoy distribution in the right tail. MS-GF+, however, can also return the score for the second best candidate PSM (rank 2), i.e. the peptide sequence with a theoretical spectrum that matches the observed spectrum with the second best score. The middle panel of Figure 2 shows a similar plot, but only including rank 2 PSMs. If the decoy PSM scores are a good approximation of target PSM scores, we expect the distributions of rank 2 target and rank 2 decoy PSMs to be very similar because the majority of the rank 2 target PSMs are likely to be incorrect ^18^. As expected, both distributions look very similar and are located at low values, which indicates that the majority of the theoretical spectra of the rank 2 PSMs does not match very well to their observed spectrum. Nevertheless, there are slightly more rank 2 target PSMs than rank 2 decoy PSMs. Indeed, some of the rank 2 target PSMS can be expected to be correct e.g. because their rank 1 PSM was incorrect, because of homologous sequences, or because of a chimeric spectrum, where two different peptides were cofragmented. This can also explain the few higher-scoring rank 2 target PSM. Even though rank 2 PSMs can differ in distribution from rank 1 incorrect target PSMs, the direct comparison of the distribution of rank 2 targets and decoys can in some cases help in the evaluation of decoy PSMs. Indeed, they give a better view on the comparison in the right tail of the decoy distribution, for which the full rank 1 target distribution cannot be used.

The right panel of Figure 2 is a similar plot, but here the *Pyrococcus furiosus* MS run was searched with MS-GF+ against the Swiss-Prot Human database using the competitive TDA method. As *Pyrococcus furiosus* is evolutionarily very distinct from *Homo sapiens,* none of the observed spectra should have a proper match in the database upon removal of PSMs that match to common contaminants. Again, we expect the score distributions of the targets and decoys to be very similar and located at low score values, which is indeed the case.

In the method section we showed that the TDA depends critically on two assumptions: 1) the distribution of decoy PSM scores is a good approximation of that of incorrect target PSMs, and 2) it is equally likely that an incorrect PSM matches to a decoy as to a target sequence in the concatenated database. The TargetDecoy package provides diagnostic plots to critically assess these assumptions. Note that this has to be done on *all* target and decoy PSMs, so prior to any thresholding! We therefore call upon mass spectrometrists and data analysts to always use search settings that return all PSMs and to threshold these only after assessing the TDA assumptions.

### Diagnostic plots

In the methods section we showed that the competitive target decoy approach implicitly estimates the cumulative distribution of the targets *F_t_*(*x*) using the empirical cumulative distribution function (ECDF) of all target PSM scores, *ECDF_t_*(*x*), the cumulative distribution of incorrect targets *F*_0_(*x*) using the empirical cumulative distribution function of the decoy PSM scores, *ECDF_d_*(*x*), and the fraction of incorrect target PSMs, *π*_0_, as the ratio of the number of decoy PSMs on the number of target PSMs. We further showed that the TDA-FDR requires that 1 – *ECDF_d_*(*x*) is a good approximation of 1 – *F*_0_(*x*), which is equivalent to assessing that *ECDF_d_*(*t*) is a good approximation of *F*_0_(*t*). The latter could be assessed graphically with diagnostic percentile percentile (P-P) plots if we would have a sample of *F*_0_(*t*). Indeed, P-P plots plot the ECDFs of two distributions against each other, i.e. the *ECDF*_0_(*x*) against the *ECDF_d_*(*x*) for all scores *x* that are observed, which would follow the 45-degree line if both distributions are equal. Note that we could make this plot for the *P. furiosus* search against the Swiss-Prot Human database upon removal of PSMs matching to known contaminants. Indeed none of the observed *P. furiosus* spectra have their matching sequence in the database and the cumulative distribution of the target PSMs *F_t_*(*x*) = *F*_0_ (*x*) as all target PSMs are incorrect. The dots in the P-P plots for the *P. furiosus* search against Swiss-Prot Human follow the 45-degree line through the origin, indicating that both distributions are equivalent (left panel of Figure 3).

**Figure 3:**
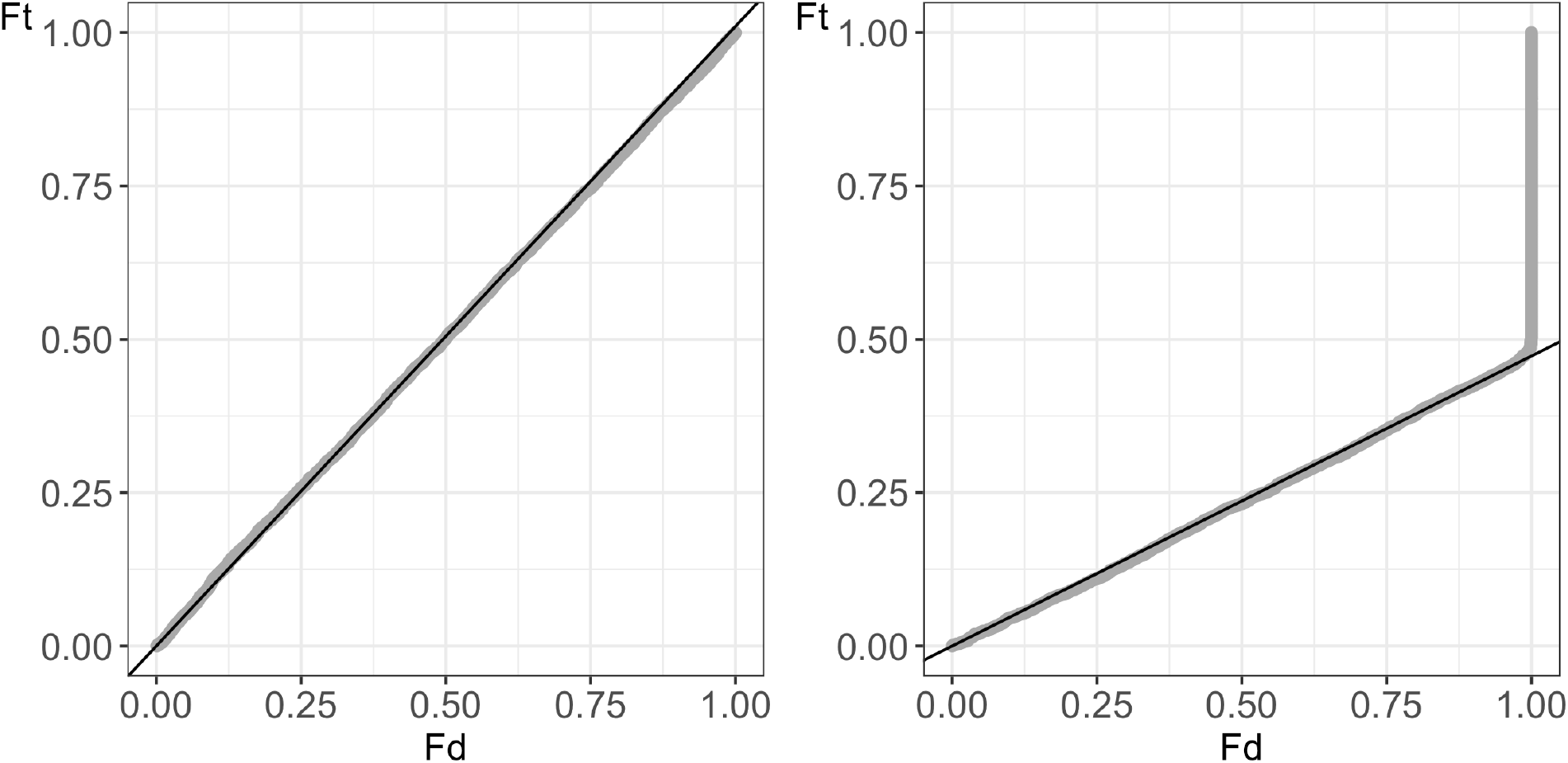
P-P plots of empirical cumulative distribution of targets against decoy for a concatenated search on a *Pyrococcus* run using MS-GF+ upon removal of PSMs matching to common contaminants. Left panel: P-P plot of target ECDF (Ft) against decoy ECDF (Fd) for a search against a human database that does not contain matches for *P. furiosus* sequences. The dots in the P-P plot follow the 45 degree line indicating that the target and decoy ECDFs are similar for all scores x. Right panel: P-P plot of target ECDF (Ft) against decoy ECDF(Fd) for search against a database with all canonical sequences for *P. furiosus* in Swiss-Prot. If the TDA assumptions are valid, we expect the P-P plot to be close to a line with a slope equal to the expected fraction of incorrect targets (*π*_0_ line) up to a certain point, then it will start to deviate from the line as higher scores are reached and correct target PSMs start to accumulate, and finally it will shoot up as the highest decoy score has been reached, and only (high scoring) target PSMs are left. The PSMs of the MS-GF+ search follow this pattern and show no deviations from the TDA assumptions.

P-P plots of the target ECDF against the decoy ECDF, however, remain very useful to evaluate the TDA assumptions in searches against the correct database, when correct target PSMs are in fact present in the search results. Indeed, for low scores the target distribution is dominated by the scores corresponding to incorrect target PSMs, hence, the distribution in the first mode of the target distribution should be very similar to that of the decoys. In this region the cumulative distribution of the targets *F_t_*(*x*) ≈ *π*_0_*F*_0_(*x*) and 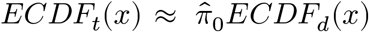. Hence the dots in the P-P plot should initially follow a straight line with slope *π*_0_ through the origin. We expect the dots to start to deviate from the *π*_0_ line when the scores increase as correct target PSMs start to accumulate at higher target scores. Eventually the dots in the plot will shoot upwards when the highest decoy score has been reached, and only (high scoring) target PSMs are left. The P-P plot in the right panel of Figure 3 for the MS-GF+ search follows this pattern and shows no deviations from the TDA assumptions.

Note, that deviations of the TDA assumptions can thus be spotted if the dots in the P-P plot do not follow a straight line at the beginning of the plot, indicating that the shape of the decoy and incorrect target PSMs differs, and/or, when the dots of the P-P plot consistently lie (below) above the *π*_0_ line. The latter indicates that target PSMs are (less) more likely to occur than decoy PSMs at all scores across the distribution, and that the assumption that an incorrect PSMs is equally likely to match to a target as to a decoy sequence is violated.

We now have two types of plots at our disposal to assess the TDA assumptions: histograms and P-P plots. Moreover, if rank 2 PSMs are present, we can use those to get a better view of how well the decoy PSM scores can approximate incorrect target PSM scores as they will also provide us with a better view on deviations in the right tail due to the drastic reduction of rank 2 target PSMs that are correct.

### Case studies

Here, we illustrate the value of our tool by evaluating the TDA for two popular search engines, MS-GF+ and X!Tandem, each used to identify peptides from a *Pyrococcus furiosus* proteome standard searched against the Swiss-Prot *Pyrococcus* database. The diagnostic plots to assess the TDA assumptions for MS-GF+ are displayed in the left panel of Figure 2 and the P-P plot in the right panel of Figure 3, and, we already argued that they are in line with our expectations under the TDA assumptions.

The diagnostic plots for X!Tandem are shown in Figure 4. Both the histogram and the P-P plot indicate that the target decoy assumptions are questionable. Indeed, the histogram shows that the shape of the decoy distribution is similar to the shape of the first mode of the target distribution. But the decoy PSMs are vastly outnumbered by low-scoring target PSMs in this region. The P-P plot still shows a straight line at lower percentiles, indicating that the shape of the decoy distribution is correct, but all dots are lying above the *π*_0_ line. This shows that there are many more target PSMs than decoys PSMs already at low scores. This could occur if the decoy database is smaller in size than the target database, when an incorrect PSM is less likely to match to a decoy than to an incorrect target sequence, or when the search engine cannot discriminate between incorrect and correct target PSMs. The former was not the case for our specific search settings as we were using a reversed decoy database. If the latter happens, the excess target PSMs should follow the tail behavior that can be anticipated for based on the second mode in distribution of targets that are dominated by high scoring correct target PSMs. Here, however, the plot clearly indicates that the number of incorrect targets seems to be underestimated by the decoys, which leads to an underestimation of the FDR.

**Figure 4:**
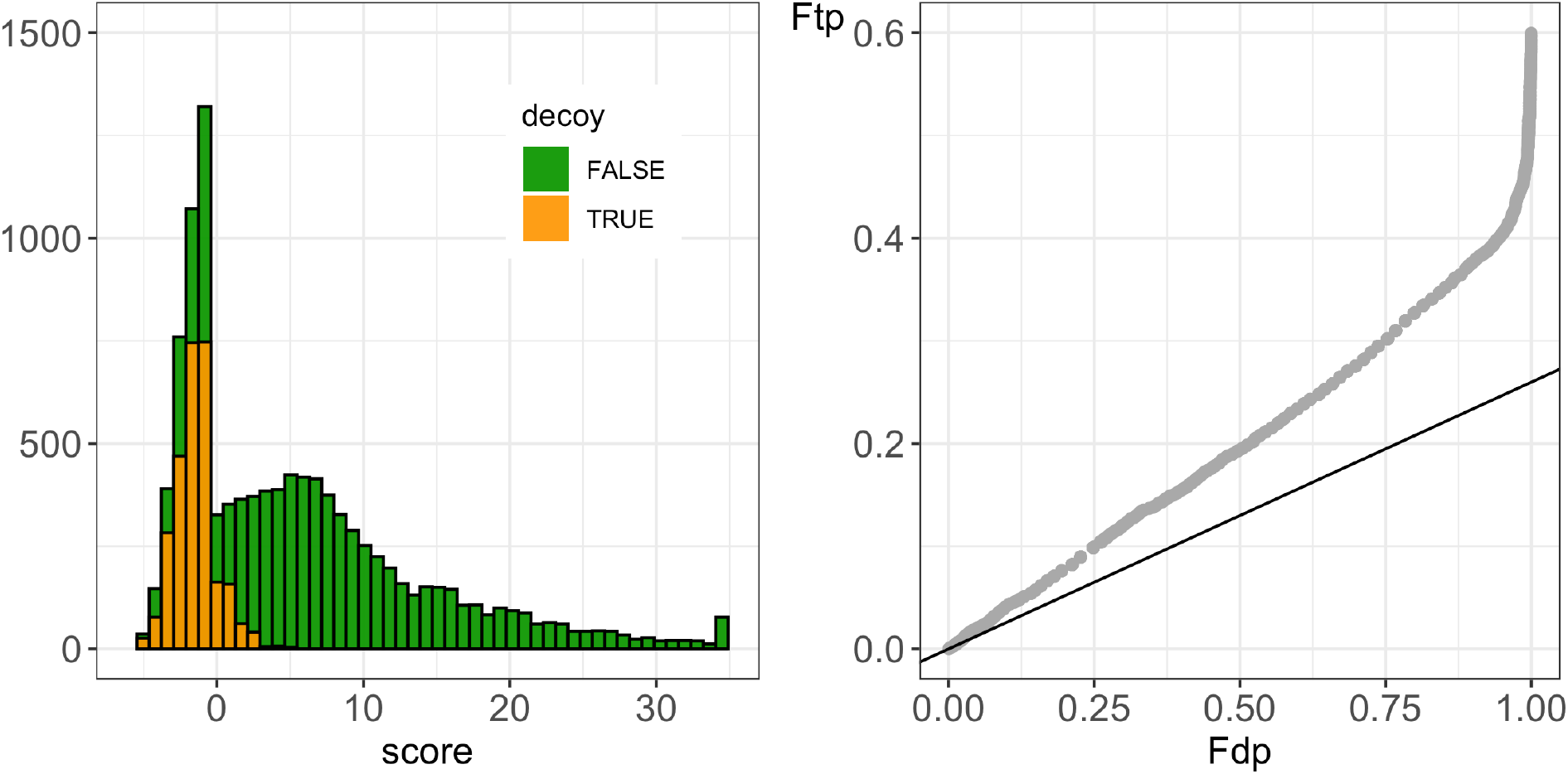
Histogram and P-P plot for a concatenated search on a *Pyrococcus* run against a database of canonical *P. furiosus* sequences from Swiss-Prot using X!Tandem with refinement. Left panel: Histogram of target and decoy PSM scores. The distribution of the target PSMs is bimodal. The shape of the decoy PSMs seems to match with the shape of the first mode of the distribution of the target PSM. However, the number of decoys is much lower than that of low scoring target PSMs. Right panel: P-P plot comparing the empirical cumulative distribution of the scores for target and decoy PSMs. The dots follow a straight line indicating that the shape of the decoy distribution is similar to that of the first mode of the target PSM score distribution. However, the dots lie systematically above the *π*_0_ line, which suggests that incorrect PSMs are more likely to match to a target than to a decoy sequence, making the TDA assumptions questionable.

We hypothesize that this is due to X!Tandems’ two-pass search strategy, which has already been reported to be a search strategy that is not TDA-compliant^3^. In the first phase, a standard search is done, which does not allow for extra modifications nor for extra missed cleavages. In a second phase, a new search is conducted solely against the identified proteins found in the first phase, but now using a more comprehensive strategy that allows for extra missed cleavages and additional post-translational modifications. To assess the impact of the refinement step, we conducted an X!Tandem search without refinement. The resulting diagnostic plots are presented in Supplementary Figure S1 and show no clear deviations from the TDA assumptions. With the results without refinement, we could classify PSMs of the search with refinement as a) novel spectra that were only matched upon refinement, b) PSMs that were matching to the same PSM, and c) PSMs that switched sequence upon refinement.

In Figure 5, we show the distributions of each of these PSM types and stratify them further according to decoys and targets. The left panel shows histograms stratified according to their identity and clearly shows that the decoys that did not switch sequence (“same decoys”) have a very similar distribution as the first mode of the distribution of targets that did not switch sequence (“same targets”) upon refinement. Moreover, the number of “same decoys” also seems to be a realistic estimate of the number of incorrect target PSMs among the targets that did not switch sequence. Upon refinement, few PSMs switch sequences and a large number of novel spectra are matched. The majority of the new PSMs match to target sequences (“new targets”) and only a few are matched to decoys (“new decoys”). The latter could be expected because the decoy space is very small during the refinement step. More importantly, it is clear that the majority of the novel target PSMs have rather low scores and the shape of their distribution is very similar to that of the decoys. Hence, the majority of the novel target PSMs are likely incorrect. As a consequence, an incorrect PSM is no longer equally likely to match to a target as to a decoy sequence, upon refinement. This is confirmed in the P-P plot in the right panel of Figure 5. Indeed, if we estimate the number of incorrect target PSMs as the sum of the number of all decoys and the number of novel target PSMs, we get a *π*_0_ line that is very close to the dots in the beginning of the P-P plot. Hence, the number of decoy matches is no longer representative for the number of incorrect target matches, which illustrates that a second pass strategy can be very detrimental for the FDR estimation when using the competitive TDA approach.

**Figure 5:**
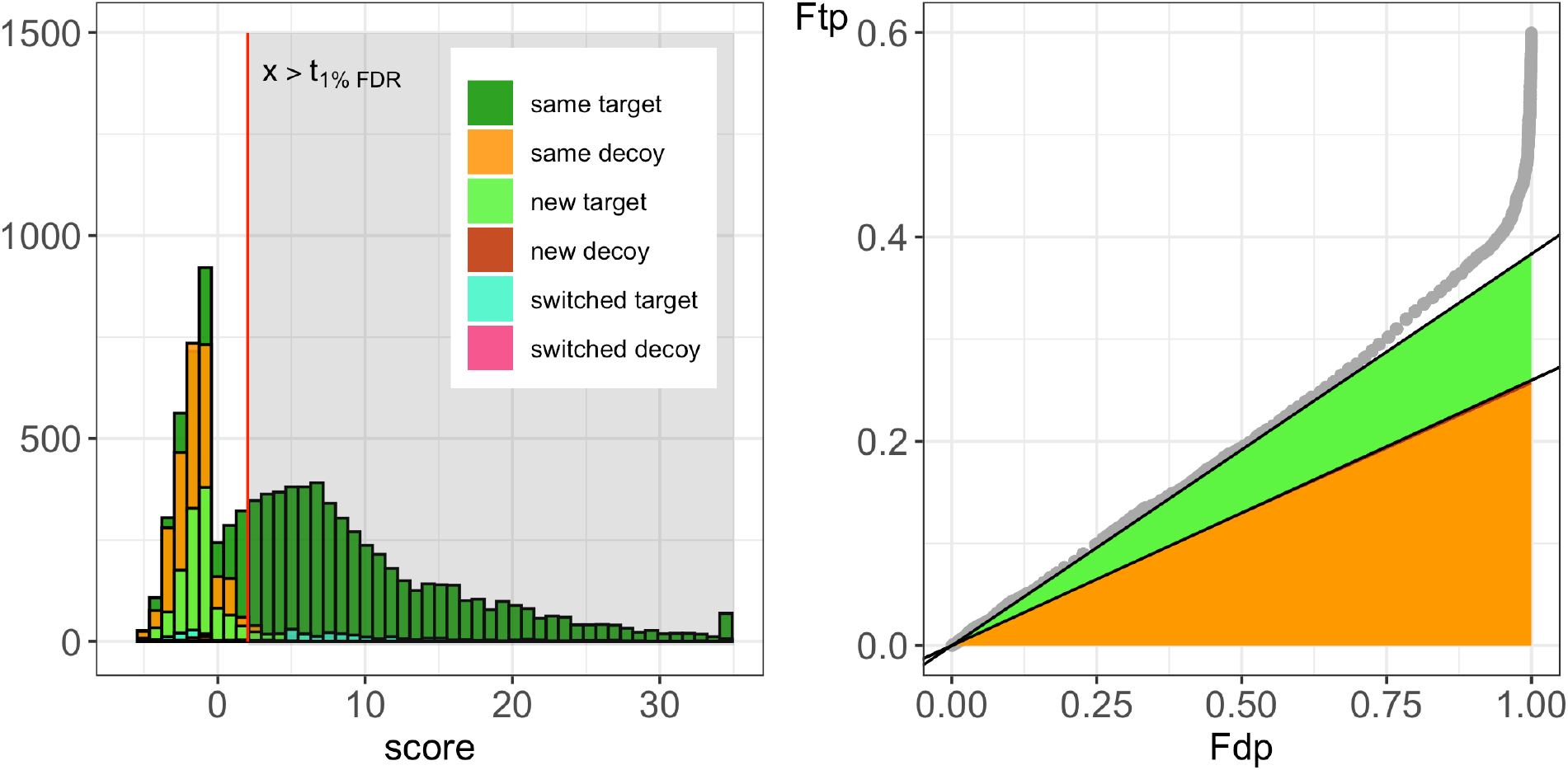
Histogram and P-P plot for a concatenated search on a *Pyrococcus* run against a database of canonical *P. furiosus* sequences from Swiss-Prot using X!Tandem while stratifying PSMs according to their match before and after refinement. Left panel: Histogram of target and decoy PSM scores upon refinement. The distribution of the target PSMs that matches to the same sequence before and after refinement (same target) is bimodal. The distribution of the decoy PSMs that match to the same decoy sequence upon refinement (same decoy) has a similar shape as the first mode of the “same target” PSM distribution. Upon refinement additional spectra can be matched (new target and new decoy). The majority of these “new PSMs” are matching to target sequences. However, their distribution is very similar to the “same decoy” distribution. Right panel: P-P plot comparing the empirical cumulative distribution of the scores for target and decoy PSMs. The ratio of the “same decoys”, “new decoys”, “switched decoys” and “new targets” on the total number of targets is represented in the plot in orange, red, pink and green. Note that there are mainly “same” decoy and “new” target PSMs, and very few “new” and “switched” decoy PSMs. When the number of incorrect target PSMs is estimated using the sum of all decoys and the number of new targets, we get a very good estimate of *π*_0_, which effectively shows that incorrect PSMs due to the refinement procedure are more likely to go to target sequences than to decoy sequences in the reduced search space of the second pass.

We also assessed the TDA assumptions for a *Homo sapiens* sample and an immunopeptidomics run searched with MS-GF+ and Andromeda, respectively. No deviations were observed that indicate a violation of the TDA assumptions (see Supplementary Figure S2 and S3). For the *H. sapiens* run searched with MS-GF+, we also had rank 2 PSMs at our disposal, which further confirms that both tails of the decoy distribution match to the distribution of incorrect target PSMs (Supplementary Figure S4). Again, slightly more rank 2 target PSMs are observed, which is not unexpected because homologous or chimeric spectra can be matched to correct sequences by rank 2 target PSMs.

Note, that diagnostic plots that do not show deviations from the TDA assumption is a necessary requirement to rely on the TDA-FDR. Do note that these plots, while useful to assess the two key assumptions of the TDA, it remains useful to explore proteomics result sets further using other metrics and visualisations to determine accuracy of the results.

Larger experiments with many runs involve multiple searches, and would require many diagnostic plots to be assessed. We therefore also provide a scaled P-P plot that allows for an efficient evaluation of the TDA assumptions for multiple searches using a single plot. This is displayed in Figure 6. In this plot, *π*_0_ is subtracted from the P-P plot for each search, allowing deviations from TDA assumptions to be spotted as deviations from the zero line, which makes comparing and combining different P-P plots straightforward. Indeed, this plot again shows no violation of the TDA assumptions for MS-GF+, and that, while the shape of the decoy distribution is correct for X!Tandem results, it seems nevertheless much more likely that a low scoring PSM matches to a target than to a decoy sequence, which will result in an underestimation of the FDR in the X!Tandem search with refinement.

**Figure 6:**
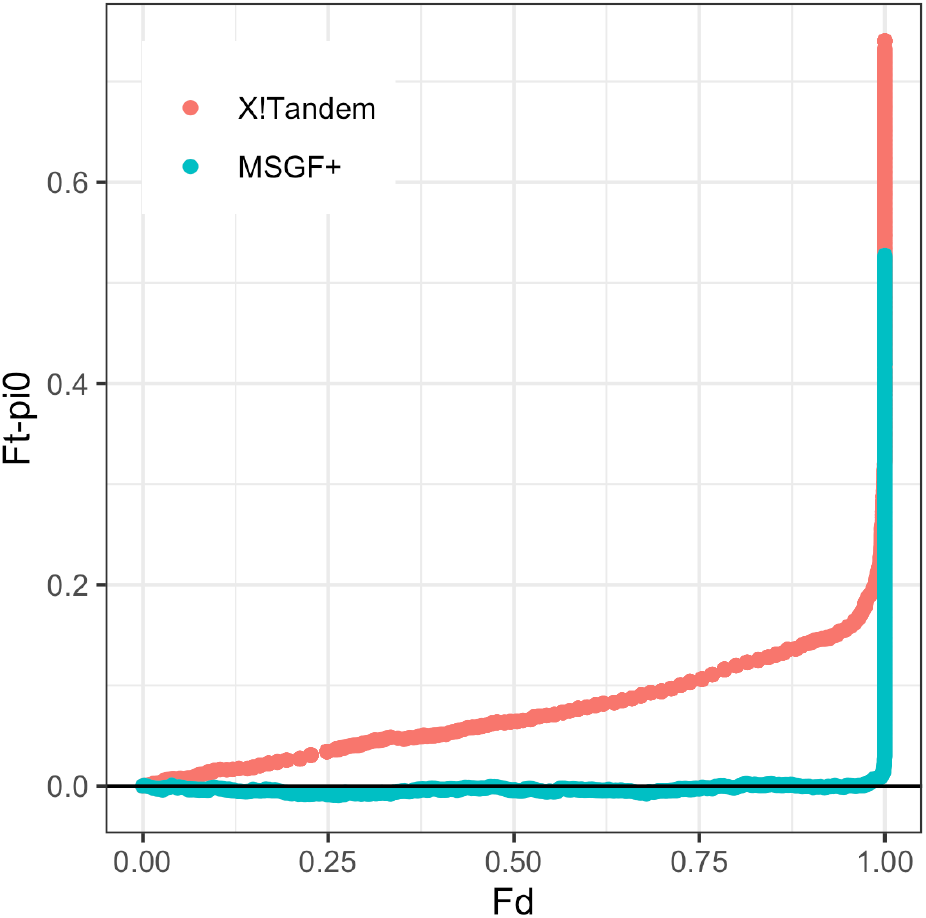
A scaled P-P plot combining P-P plots of multiple searches or runs. Here the P-P plots are combined for the *P. furiosus* run searched against canonical *P. furiosus* sequences from Swiss-Prot with MS-GF+ and X!Tandem that were also presented in the right panel of Figures 3. and 4., respectively. In the scaled P-P plot, the corresponding *π*_0_ line is subtracted from each P-P plot, so the first part of the scaled P-P plot is assumed to follow the zero line if TDA assumptions are met. The scaled P-Pplot allows one to combine search results for multiple runs and/or multiple search engines while quickly spotting for which searches the TDA assumptions are violated.

## Conclusion

The Target Decoy Approach is the default method to estimate the FDR of a set of PSMs returned after peptide identification. However, the TDA method critically relies on a number of assumptions that are typically not verified in practice. Our TargetDecoy package therefore provides a number of diagnostic plots to assess whether these TDA assumptions are met.

The provided histogram and P-P plot are very useful to evaluate the assumption that decoy PSMs provide a good simulation of target scores, and that it is equally likely that an incorrect PSM will match to a decoy or to a target sequence. Note, that the evaluation of the TDA is only possible if *all* target and decoy PSMs are available. Hence, we call upon mass spectrometrists and data analysts to always use search settings that return all PSMs and to threshold them upon assessing the TDA assumptions. For large experiments with many runs, or for comparing multiple search engines, a scaled P-P plot is provided that combines P-P plots of multiple searches in one plot, thus enabling the user to quickly spot deviating results that can then be further assessed using the default histogram and P-P plots of the TargetDecoy package.

## Acknowledgements

E.D., M.M. and L.C. are supported by Research Foundation Flanders (FWO) [G062219N]. R.G. and A.D. are supported by the Research Foundation Flanders (FWO) grants [1SE3722] and [12B7123N], respectively. L.M is supported by European Union’s Horizon 2020 Programme (H2020-INFRAIA-2018-1) [823839], Research Foundation Flanders (FWO) [G028821N], Ghent University Concerted Research Action [BOF21/GOA/033].

## Supplementary Figures

**Supplementary Figure S1:**
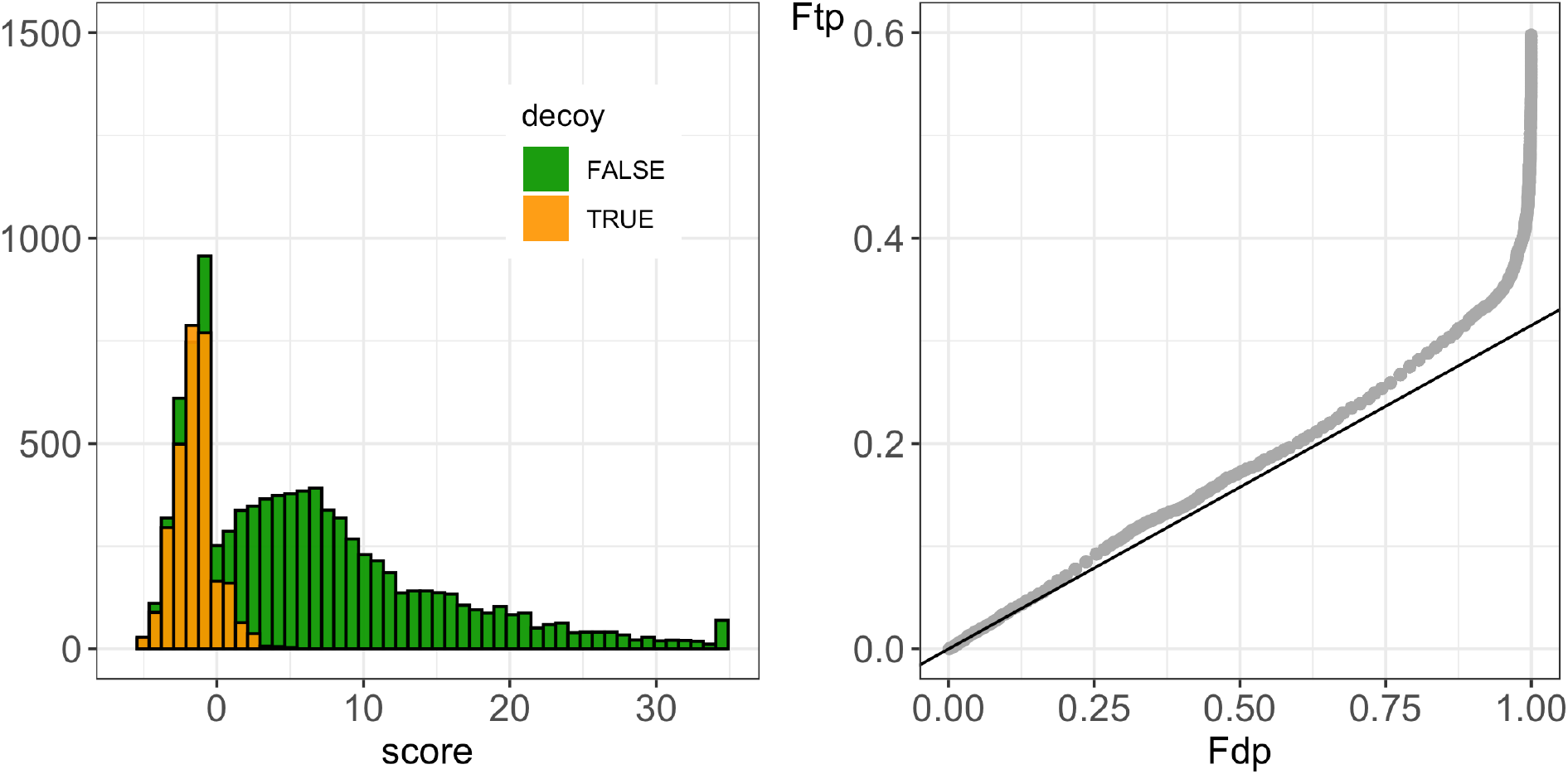
Histogram and PP-plot for a concatenated search on a *Pyrococcus* run against a database of canonical *P. furiosus* sequences from Swiss-Prot using X!Tandem without refinement. Both the histogram and the P-P plot show no violation of the TDA assumptions.

**Supplementary Figure S2:**
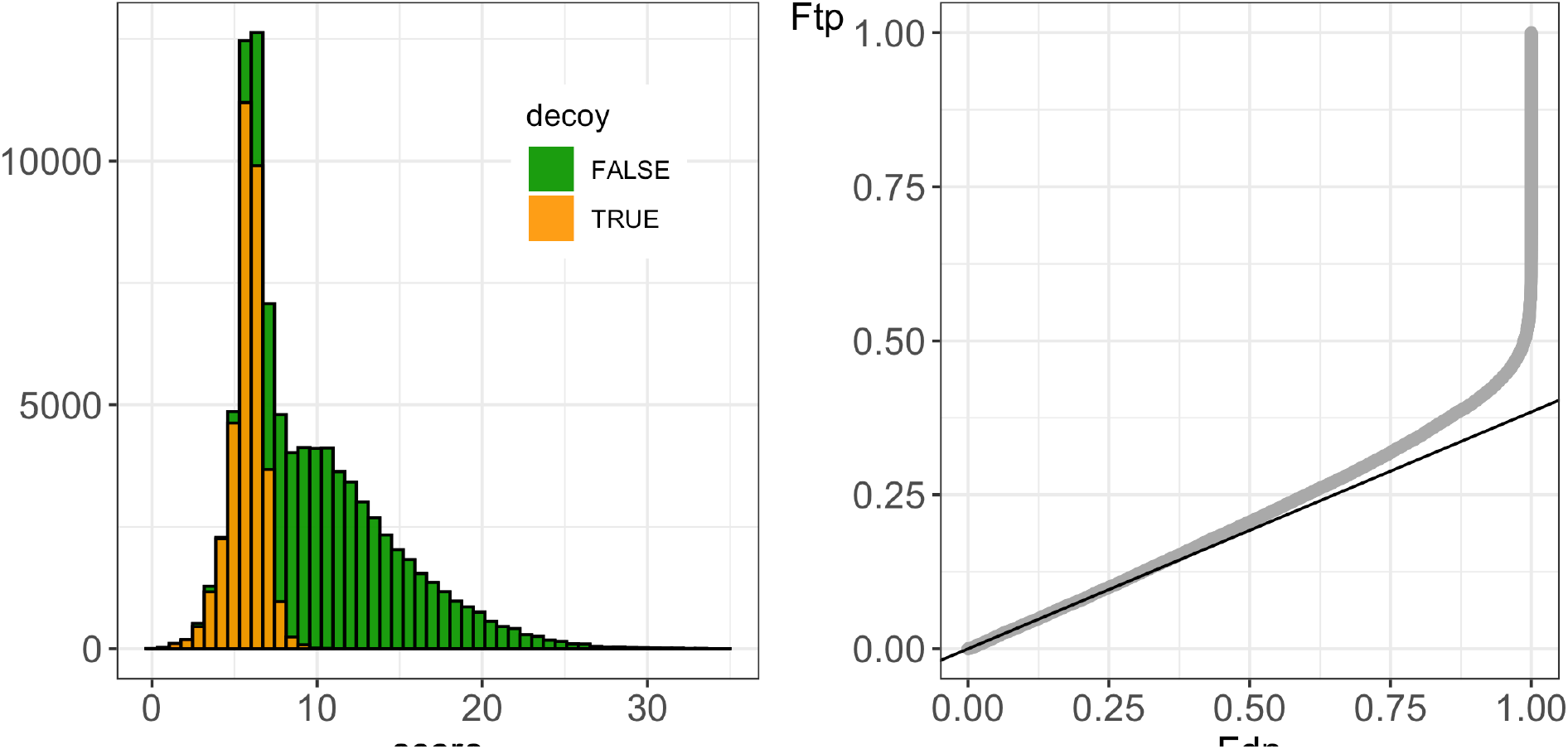
Histogram and PP-plot for a concatenated search on a *H. sapiens* run against a database of *H. Sapiens* sequences from UniProt using MS-GF+. Both the histogram and the P-P plot show no violation of the TDA assumptions.

**Supplementary Figure S3:**
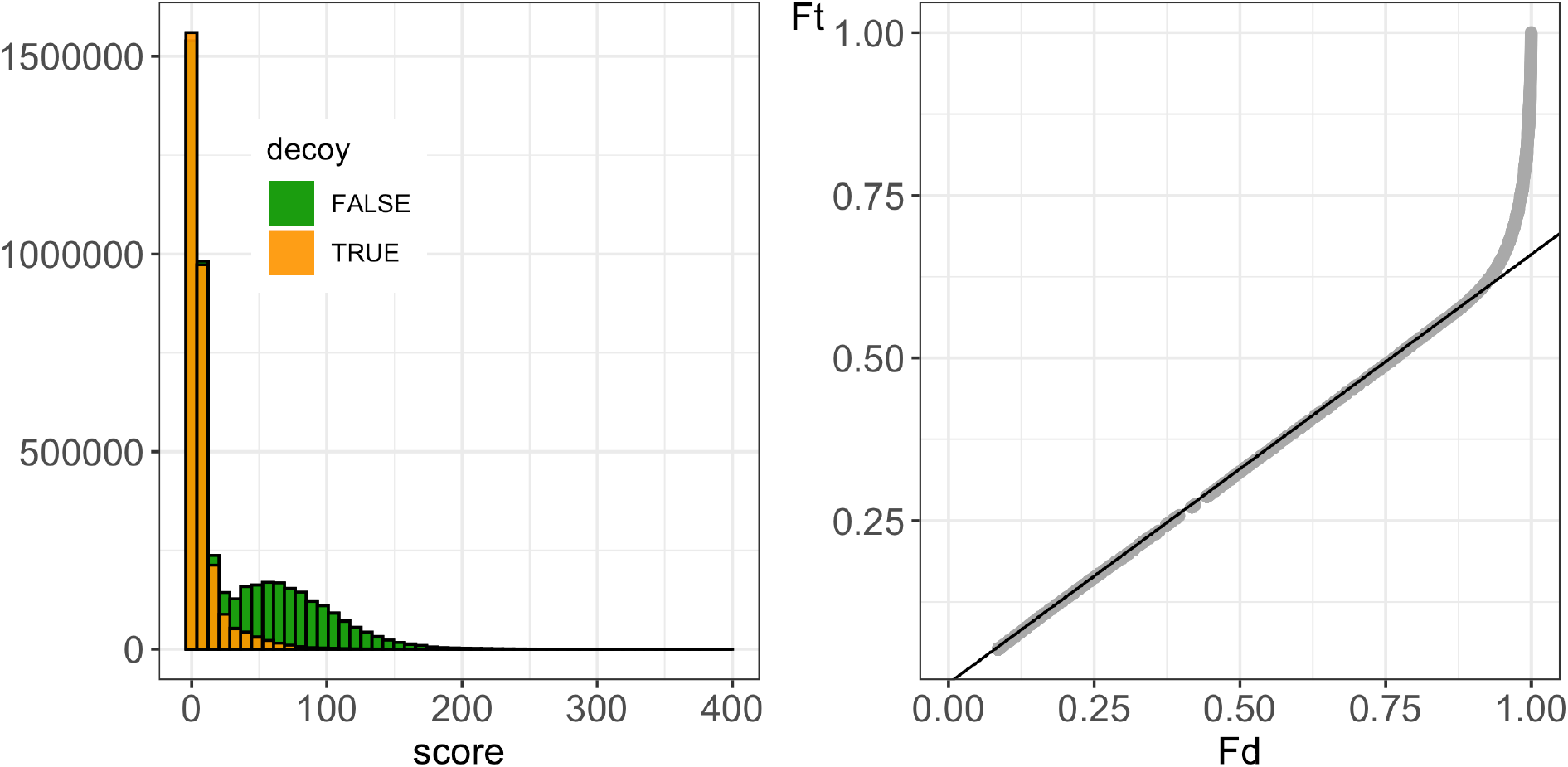
Histogram and PP-plot for a concatenated search on an immunopeptidomics run using Andromeda. Both the histogram and the P-P plot show no violation of the TDA assumptions.

**Supplementary Figure S4:**
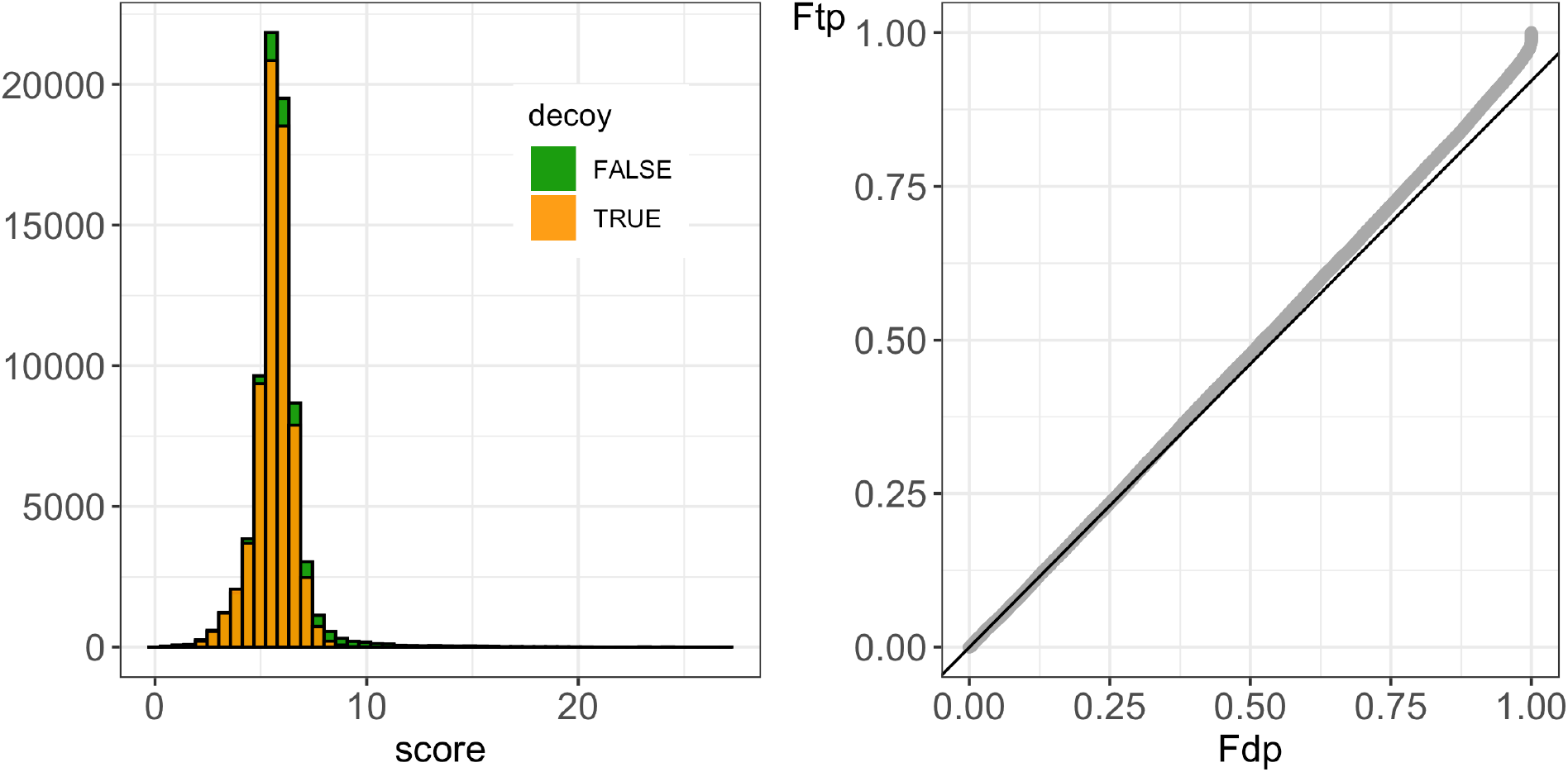
Histogram and PP-plot for rank 2 target and decoy PSM scores of a concatenated search on a H. sapiens run against a database of H. Sapiens sequences from UniProt using MS-GF+.

## Notes

https://statomics.github.io/TargetDecoyPaper/

https://doi.org/10.5281/zenodo.7308022

### Competing Interest Statement

Milan Malfait was funded by Janssen Pharmaceutical Companies of Johnson and Johnson

## References

(1) Verheggen, K.; Raeder, H.; Berven, F. S.; Martens, L.; Barsnes, H.; Vaudel, M. Anatomy and evolution of database search engines-a central component of mass spectrometry based proteomic workflows. Mass Spectrometry Reviews 2020, 39, 292–306.

(2) Elias, J. E.; Gygi, S. P. Target-decoy search strategy for mass spectrometry-based proteomics. Methods in Molecular Biology 2010, 604, 55–71.

(3) Gupta, N.; Bandeira, N.; Keich, U.; Pevzner, P. A. Target-decoy approach and false discovery rate: when things may go wrong. Journal of the American Society for Mass Spectrometry 2011, 22, 1111–1120.

(4) Efron, B. Microarrays, Empirical Bayes and the Two-Groups Model. Statistical Science 2008, 23, 1–22.

(5) Perez-Riverol, Y.; Bai, J.; Bandla, C.; García-Seisdedos, D.; Hewapathirana, S.; Kamatchinathan, S.; Kundu, D.; Prakash, A.; Frericks-Zipper, A.; Eisenacher, M.; Walzer, M.; Wang, S.; Brazma, A.; Vizcaíno, J. The PRIDE database resources in 2022: a hub for mass spectrometry-based proteomics evidences. Nucleic Acids Research 2021, 50, D543–D552.

(6) Wilhelm, M. et al. Deep learning boosts sensitivity of mass spectrometry-based immunopeptidomics. Nature Communications 2021, 12, 3346.

(7) Van Puyvelde, B. et al. A comprehensive LFQ benchmark dataset on modern day acquisition strategies in proteomics. Scientific Data 2022, 9, 126.

(8) Vaudel, M.; Burkhart, J. M.; Breiter, D.; Zahedi, R. P.; Sickmann, A.; Martens, L. A complex standard for protein identification, designed by evolution. Journal of Proteome Research 2012, 11, 5065–5071.

(9) Barsnes, H.; Vaudel, M. SearchGUI: A Highly Adaptable Common Interface for Proteomics Search and de Novo Engines. Journal of Proteome Research 2018, 17, 2552–2555.

(10) Kim, S.; Pevzner, P. A. MS-GF+ makes progress towards a universal database search tool for proteomics. Nature Communications 2014, 5, 5277.

(11) Fenyö, D.; Beavis, R. C. A method for assessing the statistical significance of mass spectrometry-based protein identifications using general scoring schemes. Analytical Chemistry 2003, 75, 768–774.

(12) Hulstaert, N.; Shofstahl, J.; Sachsenberg, T.; Walzer, M.; Barsnes, H.; Martens, L.; Perez-Riverol, Y. ThermoRawFileParser: Modular, Scalable, and Cross-Platform RAW File Conversion. Journal of Proteome Research 2020, 19, 537–542.

(13) Di Tommaso, P.; Chatzou, M.; Floden, E. W.; Barja, P. P.; Palumbo, E.; Notredame, C. Nextflow enables reproducible computational workflows. Nature Biotechnology 2017, 35, 316–319.

(14) da Veiga Leprevost, F. et al. BioContainers: an open-source and community-driven framework for software standardization. Bioinformatics 2017, 33, 2580–2582.

(15) Gabriels, R.; Declercq, A.; Bouwmeester, R.; Degroeve, S.; Martens, L. psm_utils: A high level Python API for parsing and handling peptide-spectrum-matches and proteomics search results. ChemRxiv 2022,

(16) Gatto, L.; Lilley, K. S. MSnbase-an R/Bioconductor package for isobaric tagged mass spectrometry data visualization, processing and quantitation. Bioinformatics 2012, 28, 288–289.

(17) Chang, W.; Cheng, J.; Allaire, J.; Sievert, C.; Schloerke, B.; Xie, Y.; Allen, J.; McPherson, J.; Dipert, A.; Borges, B. shiny: Web Application Framework for R. 2021; R package version 1.7.1.

(18) Gonnelli, G.; Stock, M.; Verwaeren, J.; Maddelein, D.; De Baets, B.; Martens, L.; Degroeve, S. A decoy-free approach to the identification of peptides. Journal of Proteome Research 2015, 14, 1792–1798.

